# Efficient and scalable generation of primordial germ cells in 2D culture using basement membrane extract overlay

**DOI:** 10.1101/2022.09.28.509976

**Authors:** Arend W. Overeem, Yolanda W. Chang, Ioannis Moustakas, Celine M. Roelsen, Sanne Hillenius, Talia Van Der Helm, Valérie F. Van Der Schrier, Hailiang Mei, Christian Freund, Susana M. Chuva De Sousa Lopes

**Author notes:** Corresponding author: Susana M. CHUVA DE SOUSA LOPES. Authors contributed equally (Arend W. Overeem, Yolanda W. Chang).

## Abstract

Current human primordial germ cell like cells (hPGCLCs) differentiation methods from human pluripotent stem cells (hPSCs) are inefficient, and it is challenging to generate sufficient hPGCLCs to optimize in vitro gametogenesis. We present a new differentiation method that uses diluted basement membrane extract (BME) and low BMP4 concentration to efficiently induce hPGCLC differentiation in scalable 2D cell culture. We show that BME overlay potentiated BMP/SMAD signaling, induced lumenogenesis and increased expression of key hPGCLC-progenitor markers such as TFAP2A and EOMES. These findings highlight the importance of (factors in) BME during hPGCLC differentiation, and demonstrate the potential of the BME-overlay method to interrogate the formation of PGCs and amnion in humans as well to investigate the next steps to achieve in vitro gametogenesis.

**HIGHLIGHTS:**

- hPGCLCs can be generated efficiently in 2D from hPSCs with treatment of BMP4 and BME overlay
- BME overlay method is highly scalable, cost-effective and simple to perform
- hPGCLCs differentiate together with amniotic ectoderm- and mesoderm-like cells from a TFAP2A+/CDX2+/EOMES+/GATA3+ common progenitor population
- BME overlay enables robust hPGCLC formation by potentiating BMP/SMAD signaling in the common progenitor population

## INTRODUCTION

In mammals, gametogenesis is a complex and long process that is initiated by the specification and lineage restriction of the primordial germ cells (PGCs), the founding population of the gametes (Czukiewska and Chuva de Sousa Lopes, 2022). Recapitulating (female and male) gametogenesis in vitro would enable modeling of infertility-causing diseases and may ultimately lead to new assisted reproduction techniques.

In mice, Bmp4 was identified as a crucial morphogen inducing PGC specification in the posterior-proximal epiblast (Lawson et al., 1999). Acting through the intracellular factors Smad1/5/9, Bmp4 is able to upregulate *Tbxt* (*Brachyury* or *T*) as well a specific gene regulatory network that includes *Prdm1, Prdm14* and *Tfap2c* (Saitou and Hayashi, 2021). This knowledge has led to the recapitulation of mouse PGC-like cell (PGCLC) formation in vitro, involving the exposure of mouse pluripotent stem cells (PSCs) grown as embryoid bodies (EBs) to BMP4 (Saitou and Hayashi, 2021). Subsequently, human PGCLCs (hPGCLCs) have been generated from human pluripotent stem cells (hPSCs) uncovering the divergent mechanisms of PGC specification in mice and humans as well as differences in their molecular signature (Fang et al., 2022).

The origin of PGCs in mice and humans in vivo may also differ. In mice, specified PGCs are located in the posterior-proximal epiblast at the base of the allantois shortly after the onset of gastrulation. Although it remains unknown when and where exactly PGC specification takes place in humans, in non-human primate embryos of cynomolgus monkeys PGCs were first observed in the amnion prior to gastrulation (Sasaki et al., 2016). In contrast to mice and pig that undergo amniogenesis by folding, humans and non-human primates undergo amniogenesis by cavitation (Chuva de Sousa Lopes et al., 2022; Eakin and Behringer, 2004) and may share similar origin of PGCs. In agreement, hPGCLCs share a common TFAP2A+ progenitor with amnion ectoderm-like cells in EB-differentiation assays (Chen et al., 2019) and hPGCLC formation has been demonstrated in an amniotic sac embryoid model (Zheng et al., 2019; Zheng et al., 2022).

The most widely used directed-differentiation protocols from hPSCs to hPGCLCs include aggregation into EBs and treatment with high concentration of BMP4. However, while these EB-based methods were instrumental in understanding hPGCLC formation (Irie et al., 2015; Sasaki et al., 2015), they are characterized by low efficiency and high variability per hPSC line (Chang et al., 2021; Chen et al., 2017). Reported hPGCLC yields ranged from 5%-60%, with efficiencies above 30% being uncommon, and with many hPSCs showing hPGCLC-differentiation yields below 10%. In addition, EB differentiation is low throughput, laborious, and requires harsh and stressful cell dissociation, and as a result, efficient hPGCLC generation for high throughput downstream experiments to optimize human gametogenesis in vitro. remains challenging.

In this study, we have uncovered a critical role of extracellular matrix (ECM) during hPGCLC differentiation from hPSCs in 2-dimentional (2D) culture. We show that supplementation with basement membrane extract (BME) and a concentration as low as 10ng/ml of BMP4 using an ordinary 2D cell culture format is sufficient to consistently generate high hPGCLC-yields between 30-50% within 5 days of differentiation. The hPGCLCs in this 2D system originated from a *TFAP2A+CDX2+GATA3+EOMES+* progenitor population, that also gave rise to both amniotic ectoderm-like and presumably amniotic mesoderm-like cells. Importantly, the presented hPGCLC-differentiation method is highly scalable and cost-effective, which will greatly facilitate progress achieving human in vitro gametogenesis (IVG).

## RESULTS

### Robust generation of hPGCLCs in 2D culture with BME overlay

The application of diluted BME on 2D plated hPSCs (BME overlay) has been shown to be important to induce lumenogenesis (Taniguchi et al., 2015), enabling that system to model aspects of early human embryogenesis (Karzbrun et al., 2021). As hPGCLCs formed readily in the amniotic sac embryoid model, that consisted of BMP4-treated hPSC-spheres cultured on a microfluidic device (Zheng *et al*., 2019), we hypothesized that BME-supplemented culture may facilitate the generation of hPGCLCs in regular 2D culture formats. To test this, single cell-passaged human induced PSCs (hiPSCs) were plated in mTeSR-plus medium supplemented with 2% BME (Day 0, D0) (Figure 1A). One day later (D1), the medium was replaced with previously described hPGCLC-induction medium (Kobayashi et al., 2017) containing 200ng/ml BMP4 as well as stem cell factor (SCF), leukemia inhibitory factor (LIF) and epidermal growth factor (EGF). The differentiation was carried out in hPGCLC-induction medium for 4 days (D1-D5) with the medium in the initial 2 days (D1-D3) supplemented with 2% BME (Figure 1A).

**Figure 1.**
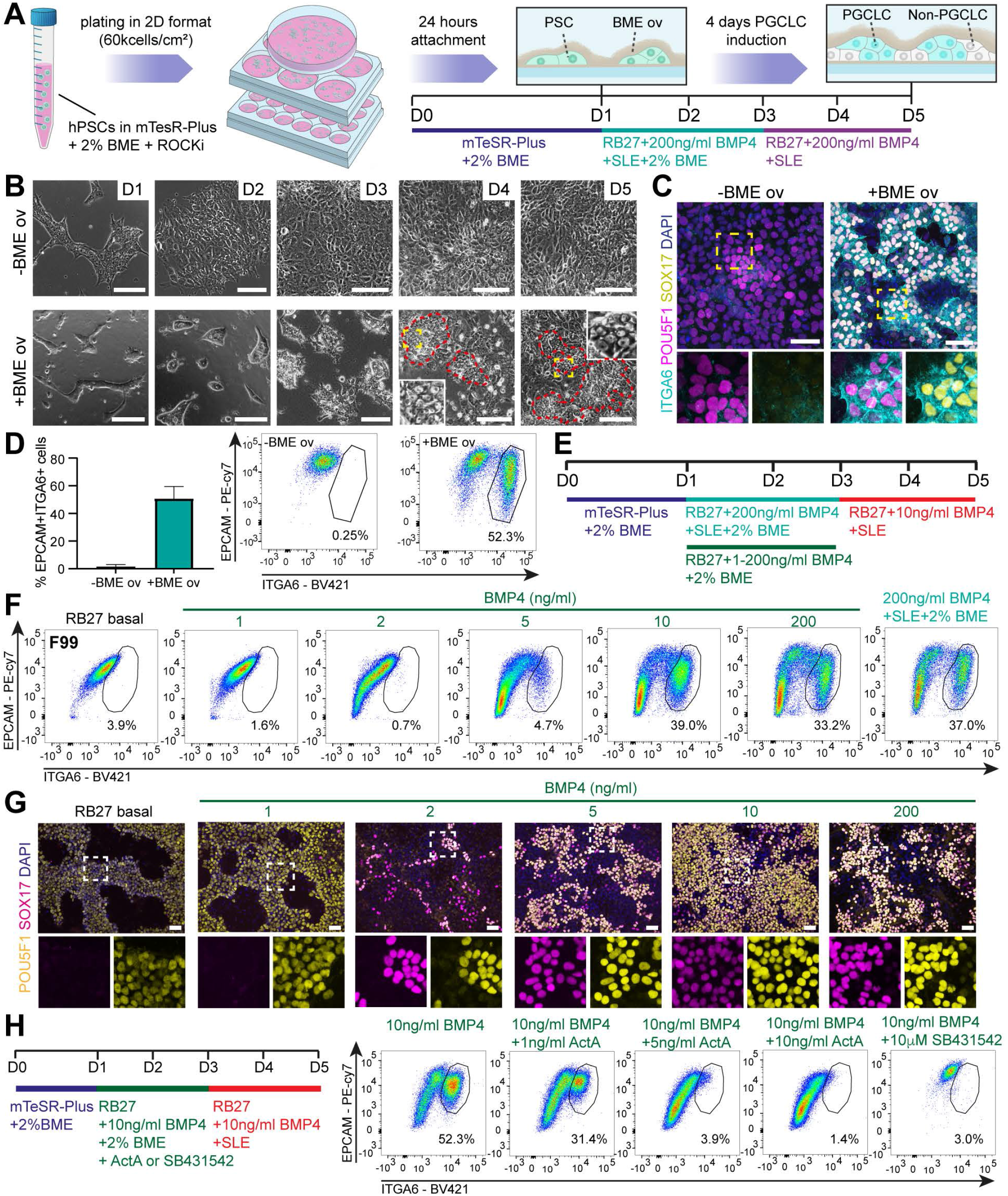
Robust generation of hPGCLCs in 2D culture using BME overlay method. **(A)** Schematic representation of the BME overlay method. Abbreviations: SLE: SCF, LIF, EGF; RB27: Advanced RPMI 1640 + B27. **(B)** Bright-field images of -BME and +BME overlay during differentiation (D1-D5). Red lines depict compacted clumps of cells. Yellow dashed box is magnified. Scale bars: 100µm. **(C)** Immunofluorescence for ITGA6, POU5F1 and SOX17 at D5 with or without BME overlay in line M54. Dashed box is magnified (bottom) showing separated channels. Scale bars: 50µm. **(D)** Bar graph (left) showing mean percentage of double EPCAM+ITGA6+ cells (n=3) at D5 with or without BME overlay in line M54 analyzed by FACS; error bars represent mean ± SD; and representative FACS plots showing the gating used (right). **(E)** Experimental scheme depicting the different conditions tested in F and G. **(F)** FACS plots depicting the percentage of double EPCAM+ITGA6+ cells at D5 in line F99 to test different BMP4 concentrations. **(G)** Immunofluorescence for POU5F1 and SOX17 at D5 in line F99 to test different BMP4 concentrations. Dashed box is magnified (bottom) showing separated channels. Scale bars: 50µm. **(H)** Experimental scheme depicting different conditions tested (left) and the associated FACS plots (right) depicting the percentage of double EPCAM+ITGA6+ cells at D5 in line F99. See also Figure S1.

Pronounced morphological differences were observed when hiPSCs were differentiated with the BME overlay (Figure 1B). In contrast to the flat colonies observed in the absence of BME overlay, tightly packed colonies were present in the presence of BME overlay. Immunofluorescence on D5 of differentiation revealed a large number of ITGA6+POU5F1+SOX17+ hPGCLCs only in the BME overlay condition across three independent hPSC lines, M54 (Figure 1C), F99 and H1 (Figure S1A), in addition to the expression of other known PGC markers, such as TFAP2C, PDPN, PRDM1 and ALPL (Figure S1B). In agreement, flow cytometry analysis using PGC markers ITGA6 and EPCAM (Mishra et al., 2021; Sasaki *et al*., 2015) revealed the generation of hPGCLCs with efficiencies of about 50% in line M54, whereas basically no hPGCLCs were detected in the absence BME overlay (Figure 1D, S1C), revealing a critical role for cell-ECM interaction during hPGCLC differentiation.

The response of hPSCs to BMP4 signaling is highly dependent on both culture format and cell density (Etoc et al., 2016; Nemashkalo et al., 2017). The hPGCLC induction medium contained a high dose of BMP4 (200ng/ml), which was optimized for EB-based methods. To establish the optimal BMP4 dosage in our 2D system, we tested different concentrations of BMP4, while removing SCF, LIF and EGF from D1-D3 and reducing the concentration of BMP4 to 10ng/ml from D3-D5 (Figure 1E). Strikingly, we observed comparable efficiencies to induce (ITGA6+EPCAM+) hPGCLCs with vastly reduced BMP4 concentrations in six independent hPSC lines (Figure 1F, S1D). Using immunofluorescence, we further confirmed an associated increase in POU5F1+SOX17+ hPGCLCs (Figure 1G).

In previous work using EB-based differentiation, we identified lines F20 and M72 as inefficient hPGCLC-generating lines (Chang *et al*., 2021). We observed the same in BME overlay differentiation, with F20 yielding 15% and M72 0.3% respectively (Figure S1D). This suggests that variance in hPGCLC generation efficiency could be an inherent cell line property and not dependent on the differentiation method used.

It was previously demonstrated that ActA/NODAL induced hPGCLC differentiation competency in hPSCs (Kobayashi *et al*., 2017). Moreover, the addition of a low dose of ActA together with BMP4 improved the specification of hPGCLCs in micropatterned colonies (Jo et al., 2022). Hence, we tested whether exogenous ActA could increase induction of hPGCLCs in our system (Figure 1H). We observed that simultaneous treatment with BMP4 and ActA from D1-D3 lowered the hPGCLCs yield in all tested concentrations, compared to treatment with BMP4 alone (Figure 1H), shortening the ActA treatment from D1-D2 gave a similarly poor outcome (Figure S1E). Interestingly, inhibiting endogenous TGFβ/ActA signalling by blocking the receptor type I (ALK4/ACVR1B, ALK5/TGFBR1, ALK7/ACVR1C) using SB431542 in our system reduced hPGCLC formation (Figure 1H, S1E).

### hPGCLCs differentiated together with amniotic ectoderm-like and mesoderm-like cells

To understand the cell types present in our BME overlay model during differentiation (2% BME overlay from D0-D3 and 10ng/ml BMP4 from D1-D5), we performed single cell transcriptomics of two hPGCLC-efficient lines (M54 and F99) and two hPGCLC-inefficient lines (F20 and M72) at D0, D2 and D5 (Figure 2A, S2A).

**Figure 2.**
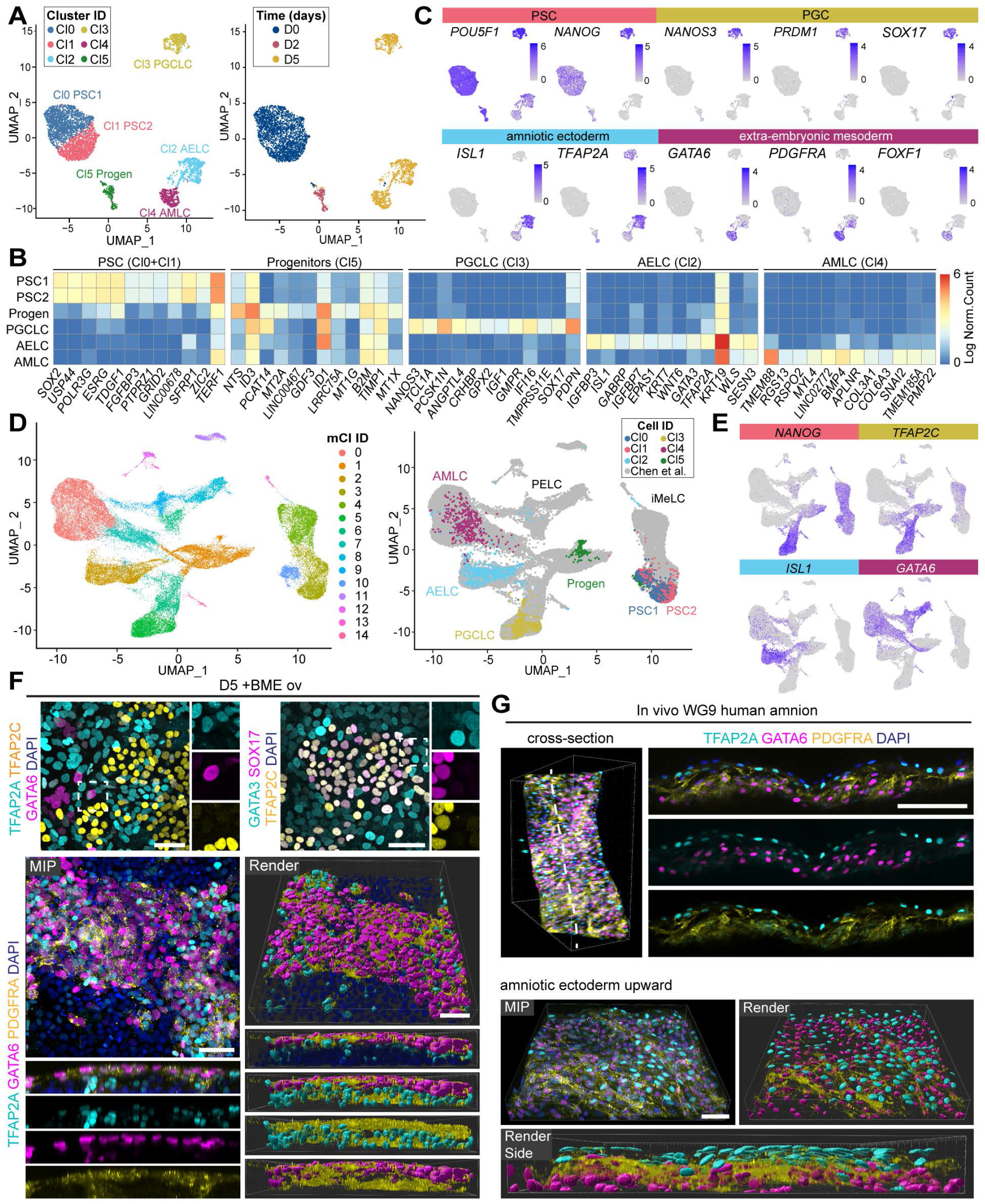
BME overlay differentiation promotes differentiation to hPGCLCs alongside amniotic ectoderm- and mesoderm-like cells. **(A)** Uniform manifold approximation and projection (UMAP) plots showing cluster identification (ID) (left) and time period in days (right) using single-cell transcriptomics of several hPSCs undergoing differentiation with BME overlay. **(B)** Heatmap showing expression levels of the top 12 DEGs of each cluster. **(C)** Expression of signature genes of cell types of interest on the UMAP plot from A. **(D)** UMAP showing integrated the single-cell transcriptomics data from EB-differentiation method (UCLA2 from Chen et al., 2019) and BME overlay method, showing overall cluster identification (left) and highlighting the cells of the BME overlay method (right). **(E)** Expression of signature genes of cell types of interest on the UMAP plot from D. **(F)** Immunofluorescence for TFAP2A, TFAP2C, GATA6 (top left), GATA3, SOX17, TFAP2C (top right) and TFAP2A, GATA6, PDGFRA (bottom) at D5 with BME overlay. In the top panels, the dashed box is magnified (right) showing separated channels. The bottom panels depict a maximum intensity projection (MIP) and render image with digital cross section showing separated channels (bottom part). Scale bars: 50µm. **(G)** Wholemount immunofluorescence for TFAP2A, GATA6, PDGFRA in human WG9 amnion. In the top panels, a digital cross section (right) shows separated channels. The bottom panels depict a MIP and render image, also showed from the side (bottom part). Scale bars: 50µm. Abbreviations: AELC, amniotic ectoderm-like cell; AMLC, amniotic mesoderm-like cell; iMeLCs, incipient mesoderm-like cells; PELC, primitive endoderm-like cell; Progen, progenitor cells. See also Figure S2.

Visualization by uniform manifold approximation and projection (UMAP) revealed the presence of six clusters (Cl0-Cl5) (Figure 2A). The top 12 most differentially expressed genes (DEGs) (based on the avg_log2FC) indicated that Cl0-Cl1 consisted of D0 hPSCs expressing high levels of *SOX2*; Cl5 consisted of D2 progenitor cells expressing high levels of BMP signaling target genes *ID1* and *ID4*; Cl3 corresponded to hPGCLCs expressing *NANOS3* and *PDPN*; Cl2 seemed to correspond to human amniotic ectoderm-like cells (hAELCs) expressing *ISL1, GATA3, TFAP2A* and *KRT7*; and Cl4 corresponded most probably to human amniotic mesodermal-like cells (hAMLCs) expressing *TMEM88, BMP4, COL3A1* and *COL6A3* (Figure 2B).

We further confirmed cell type identity by the expression of known markers genes (Chen *et al*., 2019; Chuva de Sousa Lopes *et al*., 2022; Tyser et al., 2021): hPGCLC (Cl3) and hPSCs (Cl0-Cl1) expressed high levels of *POU5F1* and *NANOG*, but *PRDM1* and *SOX17* were exclusively expressed by hPGCLCs (Figure 2C); hAELCs (Cl2) and hAMLCs (Cl4) shared high expression of *HAND1*, but only hAMLCs expressed high levels of *GATA6, PDGFRA* and *FOXF1*, whereas many cells in hAELCs expressed *ISL1, TFAP2A, VTCN1* and *IGFBP3* (Figure 2C, S2B); and a small subset of cells in Cl2 expressed key endoderm markers *FOXA2, HNF1B, HNF4A* and *SOX17* (Tyser *et al*., 2021), presumably too small to result in a separate cluster (Figure S2C). As expected, CL3 (hPGCLCs) was almost exclusively composed of cells derived from the efficient hPGCLC-generating PSCs lines M54 and F99 (Figure S2A).

To compare the developmental timeline and cell types generated using our BME overlay method with the EB-differentiation method, we merged our single-cell dataset with the single-cell dataset published by Chen and colleagues (Chen *et al*., 2019) (Figure 2D, S2D). The molecular signatures were largely similar and the small population of human endoderm-like cells (part of Cl2) now formed an independent cluster together with cells previously identified as primitive endoderm-like cells (Figure 2E, S2E). Hence, our 2D-differentiation BME overlay method is providing an alternative method for high-throughput generation of hPGCLCs, while maintaining similar cell types obtained using a 3D-differentiation method.

Next, we validated by immunofluorescence the three main cell types present at D5: TFAP2C+/SOX17+ hPGCLCs, TPAP2A+/GATA3+/SNAI2+/KRT7+ hAELCs, and GATA6+/PDGFRA+ hAMLCs (Figure 2F, S2F). Finally, using human amnion from 9 weeks of gestation (WG9), we confirmed by whole mount immunofluorescence the expression of TFAP2A in the amniotic ectoderm, and GATA6 and PDGFRA in amniotic mesoderm (Figure 2G).

### Lumenogenesis and hPGCLC differentiation were independent events

A particular feature of BME overlay culture is the formation of lumen-containing structures (Taniguchi *et al*., 2015). Since we observed distinctive morphology resembling tube/lumen structures in the BME overlay method, we investigated whether lumenogenesis is linked to the differentiation of hPGCLCs. We were able to detect laminin deposition on top of formed luminal structures on D2 differentiation with BME overlay (Figure 3A). By contrast, in the absence of the BME overlay, cells remained as a single cell layer (Figure 3A). Moreover, we observed the expression of basal-lateral markers ITGB1 and CTNNB1, apical marker PODXL and tight-junction marker TJP1 (Figure 3A, 3B, S3A), confirming lumenogenesis at D1-D2. At D3, the lumen expanded and SOX17+/PRDM1+/PDPN+ hPGCLCs were visible adjacent to the lumen (Figure 3C, 3D). At D5, the lumens lost structural integrity and a large number of SOX17+ hPGCLCs could be observed (Figure 3D, S3B). In contrast to TPAP2A+ hAELCs that expressed a clear rim of TJP1, SOX17+ hPGCLCs only showed a focal accumulation of TJP1 (Figure S3B).

**Figure 3.**
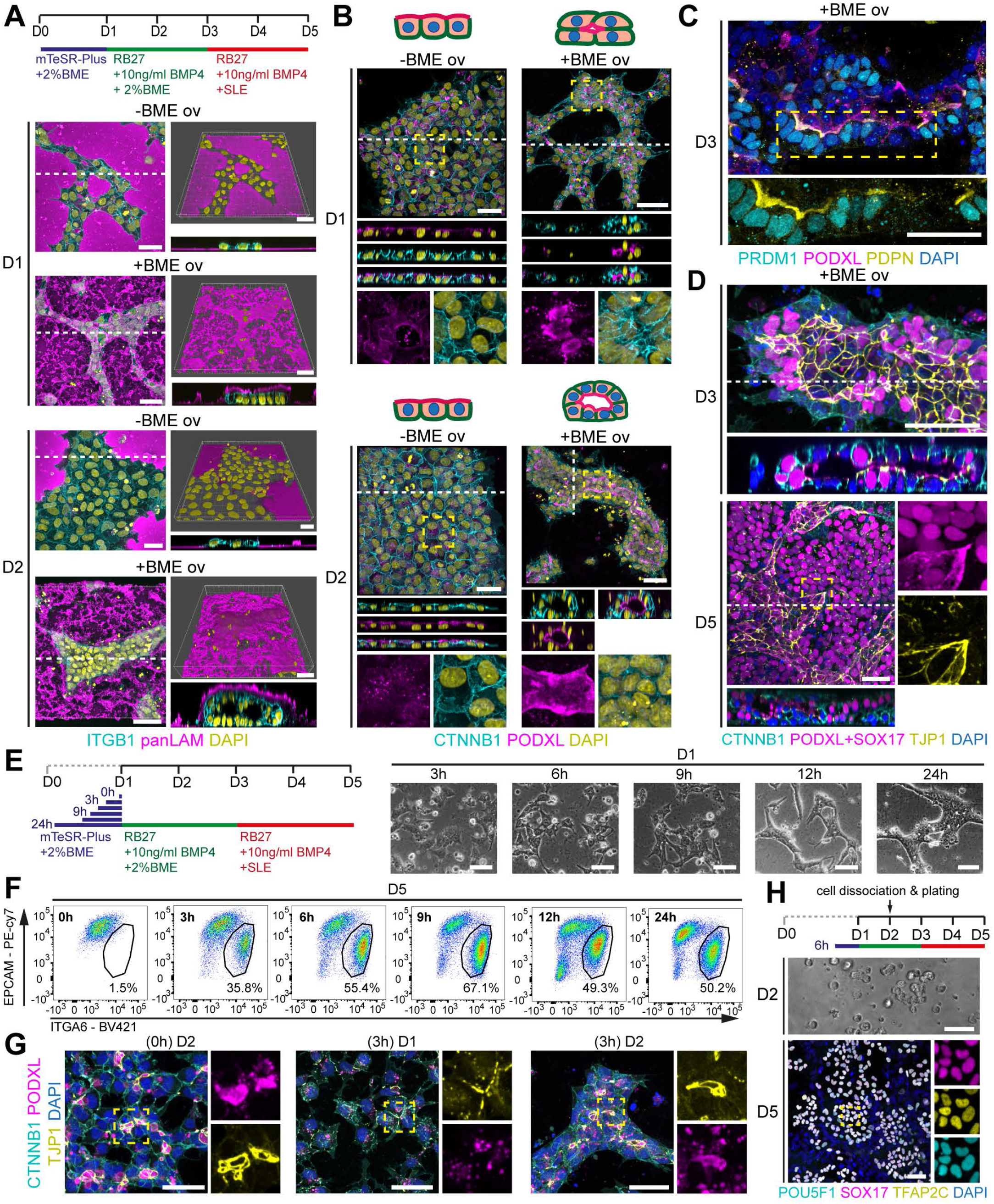
Differentiation of hPGCLCs is not coupled to lumenogenesis, but depends of BME overlay. **(A)** Experimental scheme depicting the conditions used (top) and immunofluorescence for ITGB1 and panLAM on maximum intensity projection (left) and render image (top right) at D1 and D2 with or without BME overlay. Dashed line in left panel shows the level of the digital cross section (bottom right). Scale bars: 50µm. **(B)** Immunofluorescence for CTNNB1 and PODXL at D1 and D2 with or without BME overlay. White dashed line shows the level of the digital cross section (middle panels) and yellow dashed box is magnified (bottom) showing separated channels. Scale bars: 50µm. **(C)** Immunofluorescence for PRDM1, PODXL, PDPN at D3 with BME overlay. Dashed box is magnified (bottom) without PODXL. Scale bar: 50µm. **(D)** Immunofluorescence for CTNNB1, PODXL+SOX17, TJP1 at D3 and D5 with BME overlay. White dashed line shows the level of the digital cross section (bottom) and yellow dashed box is magnified (right) showing separated channels. Scale bars: 50µm. **(E)** Experimental scheme depicting the different conditions tested in F and G (left) and associated bright-field images at D1 with BME overlay. Scale bars: 50µm. **(F)** FACS plots depicting the percentage of double EPCAM+ITGA6+ cells at D5 in line M54 to test different priming periods. **(G)** Immunofluorescence for CTNNB1, PODXL, TJP1 at D1 and D2 to test different priming periods. Dashed box is magnified (right) showing separated channels. Scale bars: 30µm. **(H)** Experimental scheme depicting the conditions used to disrupt the lumens at D2 (top), associated bright-field image at D2 after dissociation and immunofluorescence for POU5F1, SOX17, TFAP2C at D5. Dashed box is magnified (right) showing separated channels. Scale bars: 50µm. See also Figure S3.

Next, we varied the period of the initial plating step (mTesR-Plus + 2% BME) from 24 hours (h) to 0h (cells plated directly in RB27 + 10ng/ml BMP4 + 2% BME) (Figure 3E) to investigate whether hPGCLC differentiation depended on the timing of lumen formation. Although the initial plating step was necessary to obtain hPGCLC differentiation, a 3h plating-step was sufficient to obtain robust differentiation to hPGCLCs using two different lines, M54 (Figure 3F) and F99 (Figure S3C). Interestingly, independent of the duration of the initial plating step (between 0h and 24h), small lumens marked by PODXL+TJP1+ apical membrane domains were observed by D2 (Figure 3G). In the absence of BME overlay, we observed cellular polarization with the formation of a clear TJP1+ apical rim, but no lumen formation (Figure S3D).

To further test whether lumen maintenance at D2 was necessary for hPGCLCs differentiation, we disrupted the lumens at D2 by dissociating and replating the cells, followed by analysis at D5 (Figure 3H). Despite the disruption of lumens at D2, immunofluorescence revealed formation of TFAP2C+/SOX17+/POU5F1+ hPGCLCs (Figure 3H). In conclusion, lumenogenesis and hPGCLC differentiation appeared to be two independent events, and only exposure to BME (for a period as short as 3h) prior to BMP4+BME treatment (for two days) was essential for hPGCLC differentiation.

### BME overlay potentiates BMP4 signaling in the PGCLC progenitors at D2

We performed differential analysis between the PSCs at D0 (Cl0 and Cl1) and the progenitor population at D2 (Cl5) and observed that from D0 to D2 *DPPA4* and *SOX2* were downregulated, whereas many BMP responsive genes such as *ID1, ID4, GATA3, TFAP2A* and *MSX2* were upregulated (Figure 4A). In addition, TFAP2A was expressed in the progenitor cells at D2-D3, but not in PRDM1+/SOX17+ hPGCLCs at D5 (Figure 4B, S4A). Interestingly, in the absence of BME overlay, TFAP2A was basically absent at D2 (Figure S4B), whereas GATA3 showed comparable levels with or without BME overlay at D2-D3 (Figure 4C, S4C).

**Figure 4.**
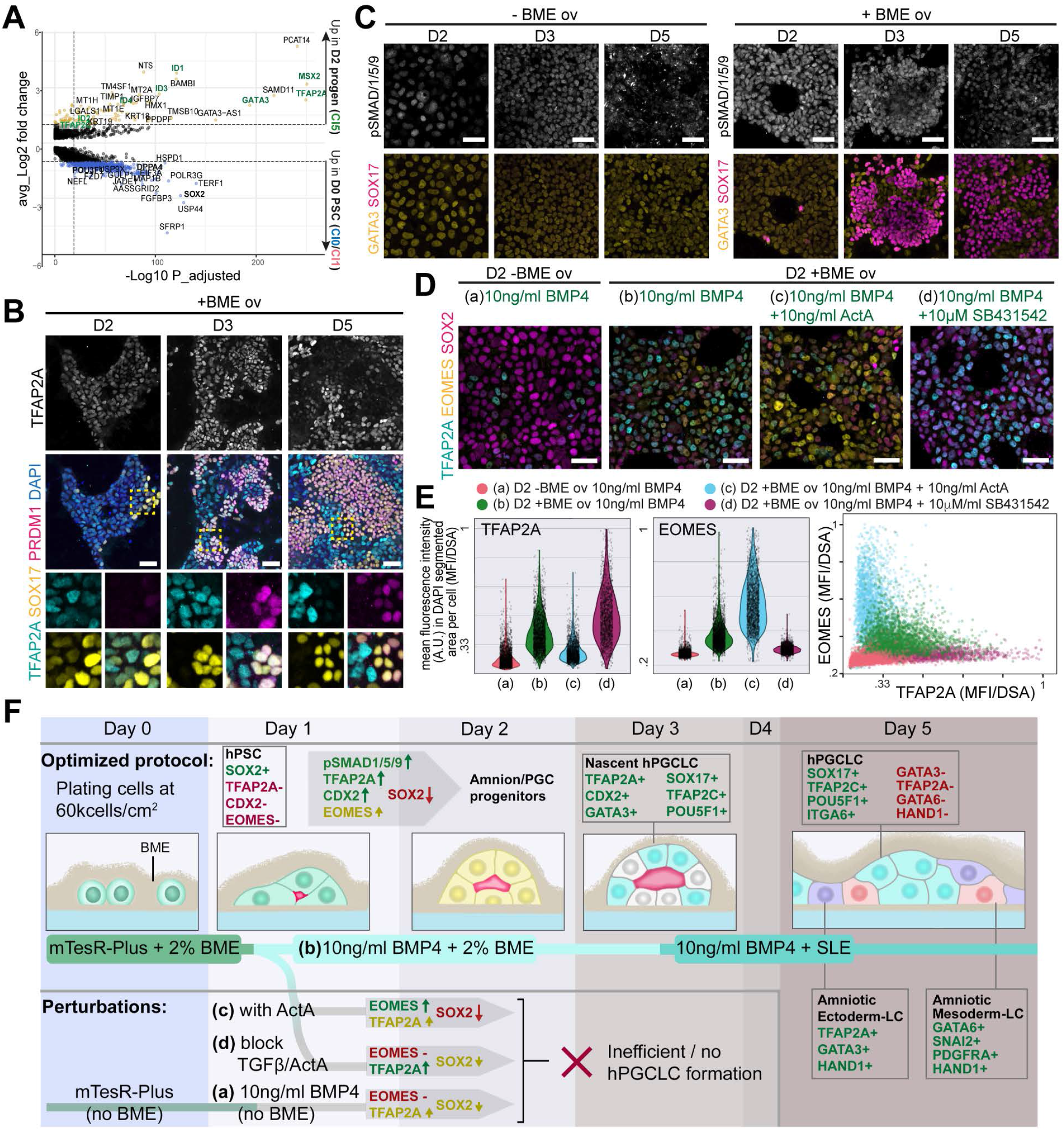
BME overlay potentiates BMP4 signaling and increases expression of critical PGC specification factors. **(A)** Volcano plot showing DEGs between hPSCs at D0 and D2-differentiated progenitors with BME overlay. **(B)** Immunofluorescence for TFAP2A, SOX17, PRDM1 at D2, D3 and D5 with BME overlay in line M54. TFAP2A is showed on top as single channel. Dashed box is magnified (bellow) showing separated channels. Scale bars: 50µm. **(C)** Immunofluorescence for pSMAD1/5/9, GATA3 and SOX17 at D2, D3 and D5 with or without BME overlay in line M54. pSMAD1/5/9 is showed on top as single channel. Scale bars: 50µm. **(D)** Immunofluorescence for TFAP2A, EOMES, SOX2 at D2 with or without BME overlay in line F99. Scale bars: 50µm. **(E)** Violin plots depict the quantification of the images in D as the mean fluorescence intensity in arbitrary units (A.U.) of TFAP2A (left) and EOMES (middle) in DAPI segmented areas (normalized to 1) per cell. The correlation between these two values per cell per condition was visualized in a scatter plot (right). **(F)** Cartoon summarizing the hPGCLC differentiation progression in the BME overlay method as well as the perturbations tested, with key analyzed markers depicted. See also Figure S4.

To test whether the BMP4 signaling activity was influenced by BME overlay, we examined the levels of phosphorylated (p)SMAD1/5/9 (Figure 4C, S4C). Even though both culture conditions (with and without BME overlay) contained 10ng/ml BMP4, the fluorescence intensity of nuclear pSMAD1/5/9 was higher in the presence of BME overlay in particular at D2 (Figure 4C, S4C).

In addition to *TFAP2A* and *GATA3, CDX2* and *EOMES* were also identified as markers of hPGCLC progenitors in EB-differentiation (Chen *et al*., 2019) and BME overlay method (Figure S4D). In agreement, similarly to TFAP2A, both CDX2 and EOMES were upregulated at D2-D3 only in the presence of BME overlay (Figure 4D, S4E, S4F). *EOMES* was previously shown to be activated by ActA/NODAL signaling during hPGCLC differentiation and to essential for initiating the hPGCLC transcriptional network (Kobayashi *et al*., 2017; Kojima et al., 2017). However, we observed that the addition of exogenous ActA was detrimental for hPGCLC differentiation in the BME overlay method (Figure 1H). To understand this discrepancy, we quantified the expression of EOMES, TFAP2A and SOX2 in the common progenitor population at D2 in the presence or absence of BME overlay; and after treatment with 10ng/ml ActA or inhibition of endogenous TGFβ/ActA signaling using 10µM SB431452 (Figure 4D).

Compared to the absence of BME overlay, the D2 progenitors cultured with BME overlay upregulated both TFAP2A and EOMES and downregulated SOX2 in line F99 (Figure 4D, 4E) and other lines F20, M72 and F31 (Figure S4F). When treated with 10ng/ml of BMP4 and 10ng/ml ActA in the presence of BME overlay, D2 progenitors upregulated EOMES considerably, whereas inhibition of endogenous TGFβ/ActA signaling blocked EOMES expression (Figure 4D, 4E), indicating that EOMES is strongly regulated by ActA signaling in our culture system. Interestingly, treatment with a combination of BMP4 and ActA from D1-D3 with BME overlay resulted at D5 in the induction of SOX17+FOXA2+ cells (Figure S4G), presumably endoderm, which is consistent with the role of ActA and its target EOMES in endoderm differentiation (Heslop et al., 2022; Yoney et al., 2022).

In conclusion, in our optimized hPGCLC differentiation method, the addition of BME overlay between D0-D3 resulted in faster downregulation of SOX2, increased pSMAD1/5/9 signaling, lumenogenesis and increased expression of TFAP2A, CDX2 and EOMES (Figure 4F). This led to the formation at D3 of nascent hPGCLCs, that downregulated TFAP2A and CDX2, upregulated *NANOS3* by D5, and which make up for about 50% of the cells in culture alongside amniotic ectoderm- and mesoderm-like cells (Figure 4F).

## DISCUSSION

The current methods to generate hPGCLCs in vitro have drawbacks regarding efficiency and scalability. As a consequence, progress regarding the optimization of protocols to further differentiate hPGCLCs into more mature germ cells, undergoing meiotic progression in vitro, both in male and in females, has been hampered. We report a new hPGCLC differentiation method that is efficient, simple, and cost effective in a highly scalable 2D format. This new method will contribute to accelerating the progression of human in vitro gametogenesis research.

Using the BME overlay, hPSCs differentiate into hPGCLC alongside two other main cell types, previously identified as amniotic ectoderm and (amniotic) extraembryonic mesoderm (Chen *et al*., 2019). We were able to verify by immunofluorescence that human amnion at 9WG consists of TFAP2A+ amniotic ectoderm and PDGFRA+/GATA6+ amniotic mesoderm, suggesting that the extraembryonic mesoderm-like cells were indeed amniotic mesoderm-like cells. The recent single-cell transcriptomics dataset of a single gastrulating human embryo, containing both amnion and hPGCs, is a tremendous resource for comparing in vitro differentiated cells to in vivo counterparts (Chuva de Sousa Lopes *et al*., 2022; Tyser *et al*., 2021), confirming that the markers used are suitable to identify amniotic cell types.

The application of BME overlay primes hPSCs to gain competency to efficiently differentiate to hPGCLCs. Priming hPSCs for 3h was sufficient and we report that (some component in) BME acted directly and quickly to potentiate BMP signaling via pSMAD1/5/9. The ECM components of BME may interact with BMP4 directly, such as in drosophila, where BMP4 homolog *Dpp* binds to collagen type IV which mediates BMP signaling (Wang et al., 2008). Alternatively, ECM may change the availability and activity of the BMP receptors. For example, ECM-integrin interactions reorder membrane into caveolae-rich lipid rafts domains (Kim et al., 2011), which is where BMPRI receptors are localized, affecting their activity (Bonor et al., 2012; Ehrlich, 2016; Hartung et al., 2006; Ramos et al., 2006). Finally, integrins activate a multitude of downstream pathways that could result in crosstalk with BMP/SMAD signaling (Kim *et al*., 2011).

Downstream of BMP4, BME-treated hPSCs showed increased pSMAD1/5/9, leading to upregulation of GATA3, TFAP2A, CDX2, and indirectly of EOMES. EOMES is essential for hPGCLC formation (Kobayashi *et al*., 2017; Kojima *et al*., 2017), but its continued and high expression has been shown to promote differentiation to endoderm (Heslop *et al*., 2022; Yoney *et al*., 2022). Consistent with this, EOMES is moderately expressed in D2-progenitor cells exposed to the BME overlay. Moreover, in agreement with EOMES being a direct target of TGFb/ActA signaling, the addition of ActA increases EOMES expression, resulting in a reduction of the hPGCLC yield and a shift in the differentiation to SOX17+/FOXA2+ endoderm like cells. Surprisingly, in the absence of BME, the D2-progenitors fail to upregulate EOMES. This may explain why 3D differentiation methods have relied on ActA/CHIR99021 preinduction, which results in *EOMES* expression.

The observation that BME potentiates BMP/SMAD signaling has significance beyond the field of *in vitro* gametogenesis. BMP4 is widely used in various differentiation protocols and models of early embryogenesis (Simunovic et al., 2019; Warmflash et al., 2014; Zheng *et al*., 2019; Zheng *et al*., 2022). Moreover, the presence of BME has proved beneficial for the development of somite-like structures in a mouse 3D gastruloid stem-cell model (van den Brink et al., 2020) as well as developing a peri-implantation assay for non-human primates (Yang et al., 2021). Hence, the combination of BME and treatment with BMP4 may prove beneficial to mimic *in vivo* processes more accurately in human models of early embryogenesis.

## LIMITATIONS OF THE STUDY

We established that a BME overlay method resulted in robust differentiation of hPGCLCs for the majority of the tested hPSC lines, including commonly used ESC line H1. However, line F20 and M72 were characterized by low hPGCLC yields. The mechanism for this line variability remains unclear, and users of the presented method will have to test hPSC lines for compatibility. In addition, the presented method is reliant on BME isolated from murine Engelbreth-Holm-Swarm (EHS) tumor, which is a complex mix of biologically active compounds that may influence differentiation outcome, including trace amounts of growth factors. Hence, the method presented is not chemically-defined nor clinical-grade.

## ACKNOWLEDGMENTS

We would like to thank the patient donating the human amnion material used in this study as well as the staff of the Gynaikon Clinic in Rotterdam. We thank members of the Chuva de Sousa Lopes group and V. Ramovs for fruitful discussions; and M. Bellin for providing the hiPSC line F20. This work was supported by the Dutch Research Council (VICI-2018-91819642 to AWO, YWC, CMR, and SMCSL), the European Research Council (ERC-CoG-2016-725722 OVOGROWTH to IM and SMCSL), the Dutch organization ZonMw (ZonMw PSIDER 10250022120001 to TVDH and SMCSL) and the Novo Nordisk Foundationgrant (reNEW NNF21CC0073729 to AWO, SH, CF and SMCSL).

## AUTHOR CONTRIBUTIONS

Conceptualization: AWO, YWC, SMCSL; Methodology: AWO, YWC, IM; Investigation: AWO, YWC, CMR, VFVDS, SH, TVDH, IM, CF; Formal Analysis (bioinformatics): AWO, YWC, CMR, VFVDS, SH, TVDH, IM, CF, HM, SMCSL; Writing: AWO, YWC, CMR, VFVDS, SH, TVDH, IM, CF, HM, SMCSL; Resources: SH, TVDH, HM, CF, SMCSL; Funding Acquisition: SMCSL; Supervision: HM, CF, SMCSL. All authors approved the final version of the manuscript.

## DECLARATION OF INTERESTS

The authors declare no competing interests.

## STAR METHODS

### KEY RESOURCE TABLE

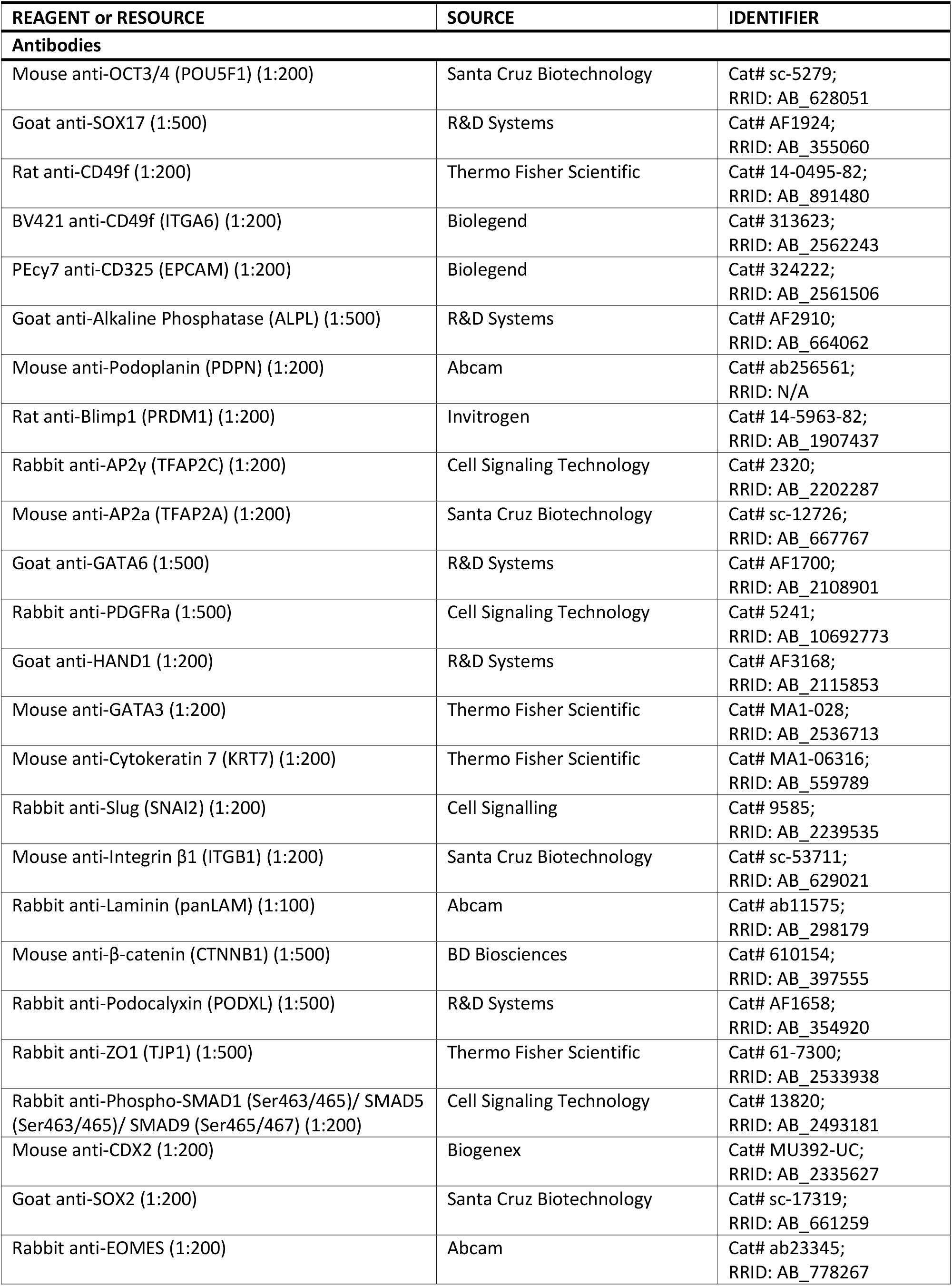
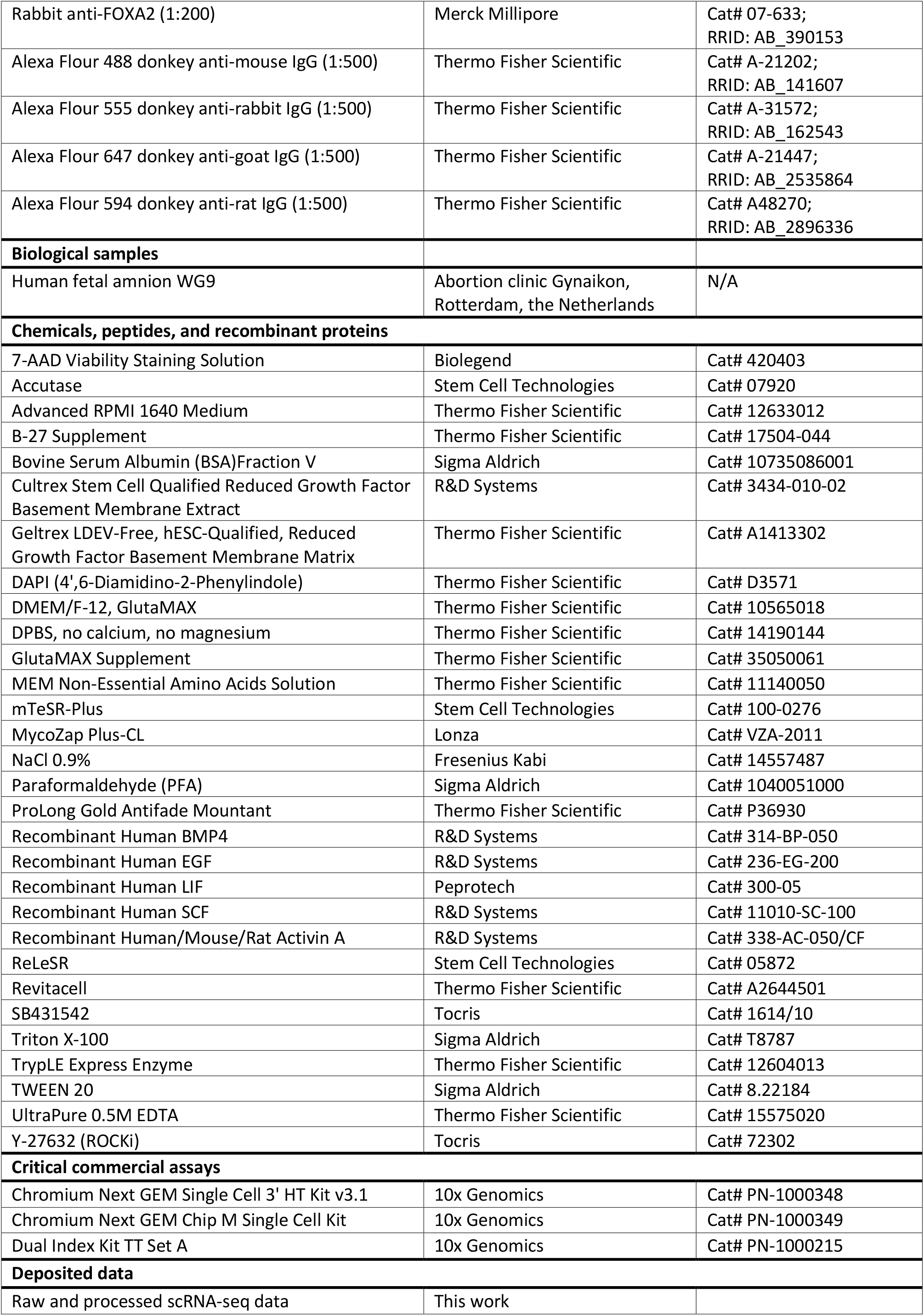
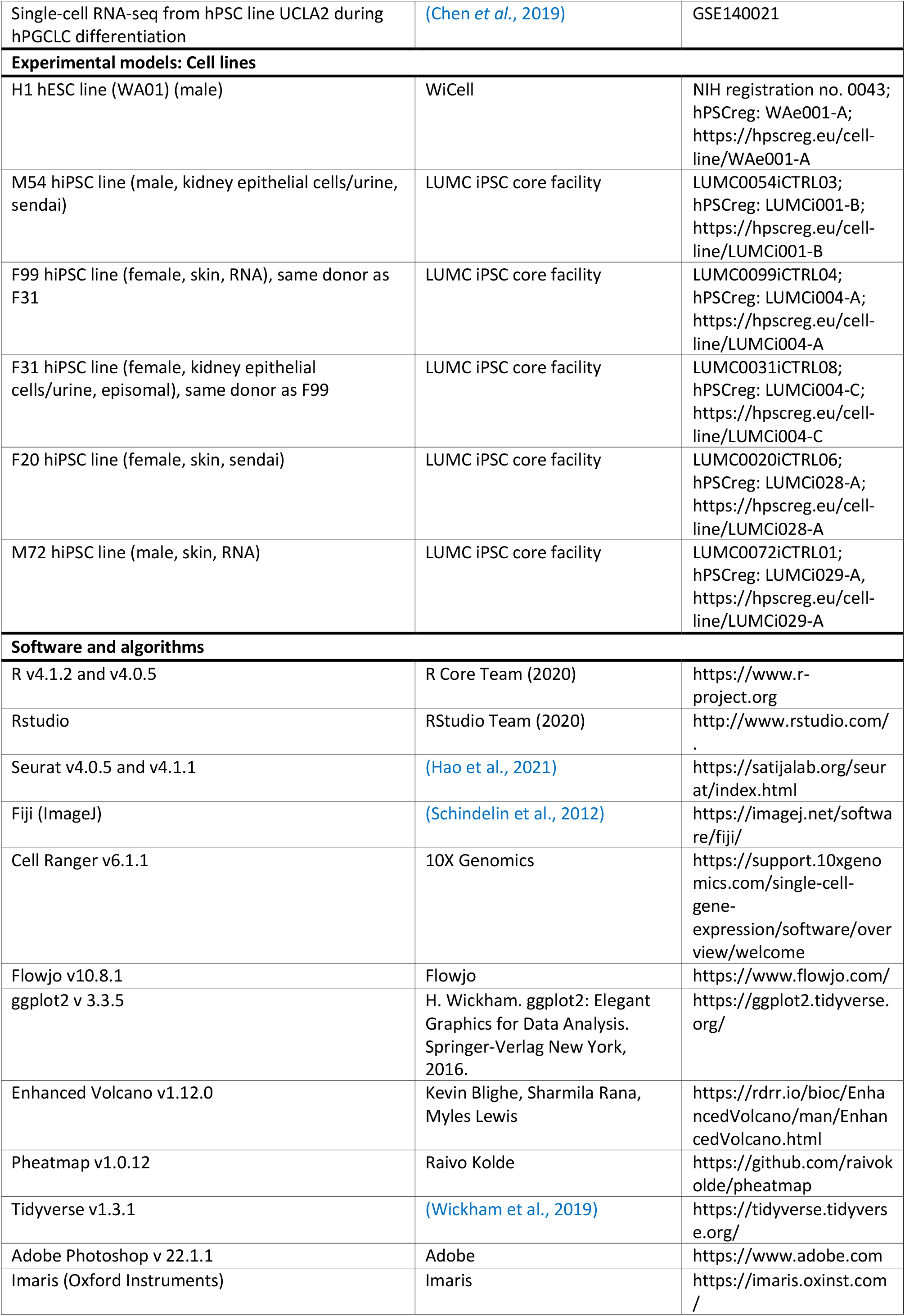
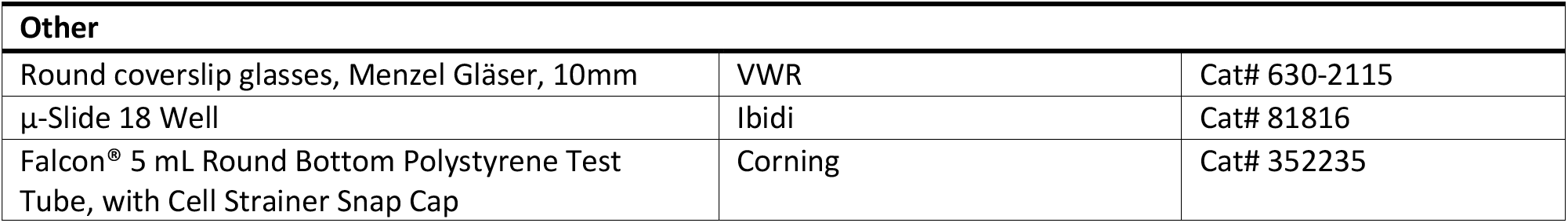

## RESOURCE AVAILABILITY

### Lead contact

Further information and requests for resources and reagents should be directed to and will be fulfilled by the lead contact Susana M. Chuva de Sousa Lopes.

### Materials availability

This study did not generate new unique materials or reagents.

## EXPERIMENTAL MODEL AND SUBJECT DETAILS

### Human samples and ethics statement

All experiments performed in this study were carried out strictly under the guidelines specified in the Declaration of Helsinki for Medical Research involving Human Subjects. For ethics approval, a letter of no objection was issued by the Medical Ethical Committee of Leiden University Medical Center (B21.054). The human amnion sample used was collected from elective abortion without medical indication, after obtaining informed consent from the donor. The amnion (2 cm x 2 cm fragment) was dissected in 0.9% NaCl solution (Fresenius Kabi), fixed in 4% paraformaldehyde (PFA) (Sigma) overnight (o/n) at 4°C washed three time in PBS, and transferred to 70% ethanol for storage at 4°C until further use.

### Routine hPSCs culture

The hPSCs used in this study were either purchased from WiCell (H1) or obtained from the LUMC hiPSC core facility (M54, M72, F20, F31, F99). All 6 lines were cultured in mTeSR-Plus media (STEMCELL Technologies) supplemented with MycoZap (Lonza) on tissue culture plates coated with either Geltrex (Thermo Fisher Scientific) or Cultrex (R&D Systems) diluted in DMEM/F12 (Thermo Fisher Scientific) at 1% (v/v) concentration. Cells were cultured at 37°C in a humidified normoxic incubator with 5% CO_2_. Routine clump passaging was performed every 4-7 days using ReLeSR (Stem Cell Technologies). The starting cultures were karyotypically normal and they were used for no more than 20 passages.

## METHOD DETAILS

### 2D hPGCLC differentiation

High quality hiPSCs of 60-80% confluency with minimal differentiation were used for hPGCLC differentiation. Briefly, cells were dissociated with TryPLE (Thermo Fisher Scientific) at 37°C for 5 minutes (min), diluted in DMEM/F12 (Thermo Fisher Scientific) to stop digestion, and spun down. Single cell suspension was resuspended in cold mTeSR-Plus media containing RevitaCell supplement (Thermo Fisher Scientific) and 2% Geltrex or Cultrex at 2.04 × 10^5^ cells/ml. We have obtained comparable hPGCLC differentiation using 10µM Y-27632 (Stem Cell Technologies) instead of RevitaCell. Depending on the plate format, the desired volume of cell suspension was added to the Geltrex- or Cultrex-coated plate to achieve a final plating density of 60,000 cells/cm^2^. On Day 1 (24 hours after plating), the medium was aspirated and the cells were washed once with aRB27 basal medium [advanced RPMI1640 (Thermo Fisher Scientific) supplemented with B27 (1:100) (Thermo Fisher Scientific), 1X Glutamax (Thermo Fisher Scientific), 1X MEM Non-Essential Amino Acids (Thermo Fisher Scientific) and Mycozap (Lonza)]. After washing, the differentiation media consisting of aRB27 with 2% Geltrex or Cultrex (+BME overlay) and 10ng/ml BMP4 (R&D Systems) was added to the cells [or with variations: omission of BME, varying BMP4 concentration and addition of SCF, LIF EGF, ActA (R&D Systems) or SB431542 (Tocris)]. Media exchange was performed the next day (D2) and on day 3 (D3), the medium was switched to aRB27 basal medium with 10 ng/ml BMP4, 10 ng/ml human LIF (PeproTech), 50 ng/ml SCF (R&D Systems) and 50 ng/ml EGF (R&D Systems). Medium change was performed daily until D5.

### Flow cytometry

Single cell suspension of the hPGCLC differentiation culture was generated by incubating the cells with Accutase (Stem Cell Technologies) for 15 min at 37°C, followed by vigorous pipetting to break up any clumps. The cell suspension was then passed through a 35µm nylon mesh strainer (Corning), pelleted by centrifugation, washed once in FACS buffer [DPBS (Thermo Fisher Scientific) with 0.5% BSA (Sigma-Aldrich)], pelleted by centrifugation and incubated with conjugated-antibodies diluted in FACS buffer at approximately 0.5-2× 10^6^ cells/ml at 4°C for 30 min. Thereafter, the cell suspension was pelleted by centrifugation and cells were resuspended in FACS buffer containing 7AAD (BioLegend, 1:100). The flow cytometry analysis was performed on an LSR-II flow cytometer (BD Biosciences). The FACS data were collected from FACSDiva Software (BD Biosciences) and FlowJo Software (BD Biosciences) were used for analysis.

### Immunofluorescence and imaging

Cells cultured for imaging were either cultured on glass coverslips (VWR) or 18-well μ-slides (Ibidi). At the time of analysis, cells were fixed in 4% PFA for 15 min at room temperature (rt). The fixed cells were then washed three times with PBS and permeabilized with 0.3% Triton-X100 (Sigma-Aldrich) diluted in PBS for 15 min at rt, followed by three washes with PBST (0.05% Tween 20 (Sigma-Aldrich) in PBS). Cells were treated with blocking buffer (1% BSA diluted in PBST) for 1 hour at rt and incubated with primary antibodies diluted in blocking buffer at 4°C o/n. Next, the cells were washed three times with PBST and incubated with secondary antibodies and DAPI (Life Technologies) diluted in blocking buffer for 1 hour at rt, followed by three PBS washes. Cells in 18-well μ-slides were imaged directly, cells on coverslips were mounted with ProLong Gold (Life Technologies). Imaging was performed on either a 200 Series Dragonfly spinning disk confocal microscope (Andor) or a TCS SP8 confocal microscope (Leica). Renderings of image stacks were generated using the Imaris surface creation functionality. Images were processed in Fiji (ImageJ2) and Adobe Photoshop (Adobe).

### Wholemount immunofluorescence and imaging

Amnion (2 cm x 2 cm fragment) was washed three times 30 min in PBS at rt to remove the 70% ethanol solution. The tissue was permeabilized with 0.5% Triton-X100 in PBS for 1 hour at rt, followed by o/n blocking in 0.2% Triton-X100 with 1% BSA in PBS (whole mount blocking buffer) at 4°C. The tissue was then incubated with primary antibodies diluted in whole mount blocking buffer for 24 hours at 4°C, washed three times in PBST for 15 min at rt and incubated with secondary antibodies diluted in whole mount blocking buffer 3 hours at rt. Finally, the tissue was washed three times with PBS for 15 min, mounted between cover glass and glass slide using Prolong Gold and imaged on a 200 Series Dragonfly spinning disk confocal microscope (Andor).

### Preparation cells for single-cell RNA-sequencing

BME-overlay cultures (D2 and D5) and hPSCs (D0) were digested with Accutase at 37 °C for 15 min or shorter if single cell suspensions were obtained. Samples were washed once in FACS buffer and centrifuged. Cells were then resuspended in FACS buffer, filtered through the FACS tube strainer cap and treated with 7AAD diluted in FACS buffer at 1:100 dilution, on ice for 3 min. Live cells were sorted on a CytoFLEX SRT benchtop cell sorter (Beckman). The collected live cells were sent to the Leiden Genome Technology Center (LGTC) for library preparation using the Chromium Next GEM Single Cell 3’ HT Kit v3.1 (10x Genomics) according to the manufacturers’ instructions and sequenced on NovaSeq6000 with V1.5 chemistry (illumina) at Genome Scan.

## QUANTIFICATION AND STATISTICAL ANALYSIS

### Primary and secondary analysis of single-cell RNA-sequencing data

Raw RNA sequencing data was processed using the Cell Ranger pipeline (v6.1.1) in which reads were aligned to the human reference genome (GRCh38) and gene UMI count matrices were generated based on gene annotation in Cell Ranger reference annotation version GRCh38-2020-A. The R package Vireo2 (v0.2.3) was used to distinguish cells from different cell lines (N=4) based on genetic variation. Count matrices generated with Cell Ranger were analyzed using Seurat (v4.1.1) workflow in R (v4.0.5). Functions mentioned below are part of the Seurat workflow unless specified otherwise. For quality control, cells expressing <2000 or >7000 genes or cells having >100000 UMIs were excluded from further analysis. In addition, cells with > 10%, or < 0.1% of the total UMIs coming from mitochondrial genes were excluded. Cells with > 6% of UMIs mapping to dissociation-induced genes were excluded as well (van den Brink et al., 2017). Data was log-normalized using the NormalizeData function (scale factor: 100000). To focus on cell type specific characteristics, batch effect between cell lines (N=4) was corrected using the fastMNN function from R package batchelor (v1.6.0). The top 2000 variable features (genes) were selected (function: FindVariableFeatures) to perform Principal Component Analysis (PCA) (function: RunPCA). The first 15 principal components (PCs) were used to calculate cell clusters, with resolution parameter set to 0.3 (functions: FindNeighbors, FindClusters). Cells were visualized on a two-dimensional plot calculated using the Uniform Manifold Approximation and Projection (UMAP) algorithm (function: RunUMAP). Differentially expressed genes (DEGs) for each cluster were calculated using function FindAllMarkers (parameters: only.pos = TRUE, min.pct > 0.25 and logfc.threshold > 0.25). Expression of individual genes were visualized using functions FeaturePlot or VlnPlot. To generate heatmap data, mean expression of each gene was calculated per cluster using base R functions, which were then filtered for top 12 differentially expressed genes per cluster (ranking based on highest fold change), and visualized using the pheatmap package (v1.0.12). For the combined analysis with Chen et al data set (UCLA2) (Chen *et al*., 2019), we applied the same filtering parameters with regard to global and mitochondrial gene expression as described above. Batch correction was done on the two replicates in the Chen et al. data (UCLA2) using the fastMNN function. The same Seurat workflow was then applied as for the in-house dataset, with changes in the following parameters: UMAP resolution parameter 0.27.

### Image quantification and visualization

The quantification of fluorescence intensity was performed using Fiji for image handling and R package ggplot2 (v3.3.5) for data visualization. First maximum intensity projections were generated on Dragonfly confocal image stacks (2048 × 2048 pixels; z-value: 10-12), for three fields of view per analyzed condition. Images were segmented based on DAPI channel to obtain areas representing nuclei. Segmentation consisted of thresholding DAPI signal, generating an image mask, and performing additional segmentation using the Fiji Watershed algorithm to separate overlapping nuclei. “Analyze particles” function was used to obtain regions of interest (minimum size set to 40 pixels), and mean fluorescence intensity signals of nuclear markers in other channels were determined in these areas. Data were loaded in R for filtering and visualization using ggplot2 (geom_jitter and geom_violin). Data was filtered for clear outliers that were a result of staining artefacts and autofluorescent debris. In addition, areas above 300 pixels which likely represented multiple nuclei were removed from the dataset.

## Notes

### Competing Interest Statement

The authors have declared no competing interest.

