## Supplementary material for "Efficient and scalable generation of primordial germ cells in 2D culture using basement membrane extract overlay": Suppl Figures S1-S4

### SUPPLEMENTAL INFORMATION

### SUPPLEMENTAL FIGURES AND FIGURE LEGENDS

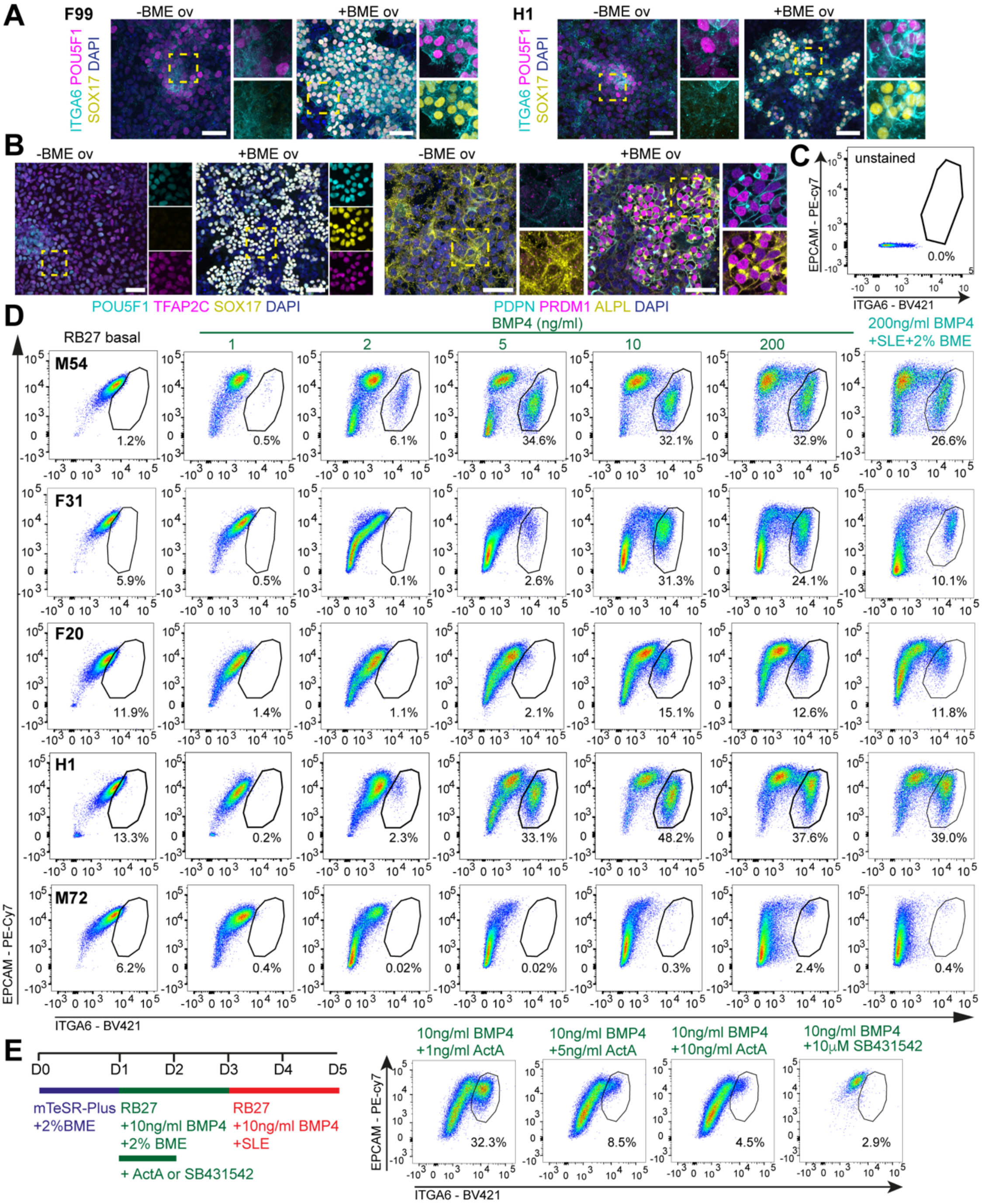


**Figure S1. hPGCLC differentiation using BME overlay method across multiple cell lines**

**(A)** Immunofluorescence for ITGA6, POU5F1 and SOX17 at D5 with or without BME overlay in line F99 and H1. Dashed box is magnified (right) showing separated channels. Scale bars: 50μm.

**(B)** Immunofluorescence for POU5F1, TFAP2C and SOX17 (left) and PDPN, PRDM1 and ALPL (right) at D5 with or without BME overlay in line M54. Dashed box is magnified (right) showing separated channels. Scale bars: 50μm.

**(C)** Unstained FACS control for Figure 1D.

**(D)** FACS plots depicting the percentage of double EPCAM+ITGA6+ cells at D5 in different lines to test different BMP4 concentrations.

**(E)** Experimental scheme depicting different conditions tested (left) and the associated FACS plots (right) depicting the percentage of double EPCAM+ITGA6+ cells at D5 in line F99.

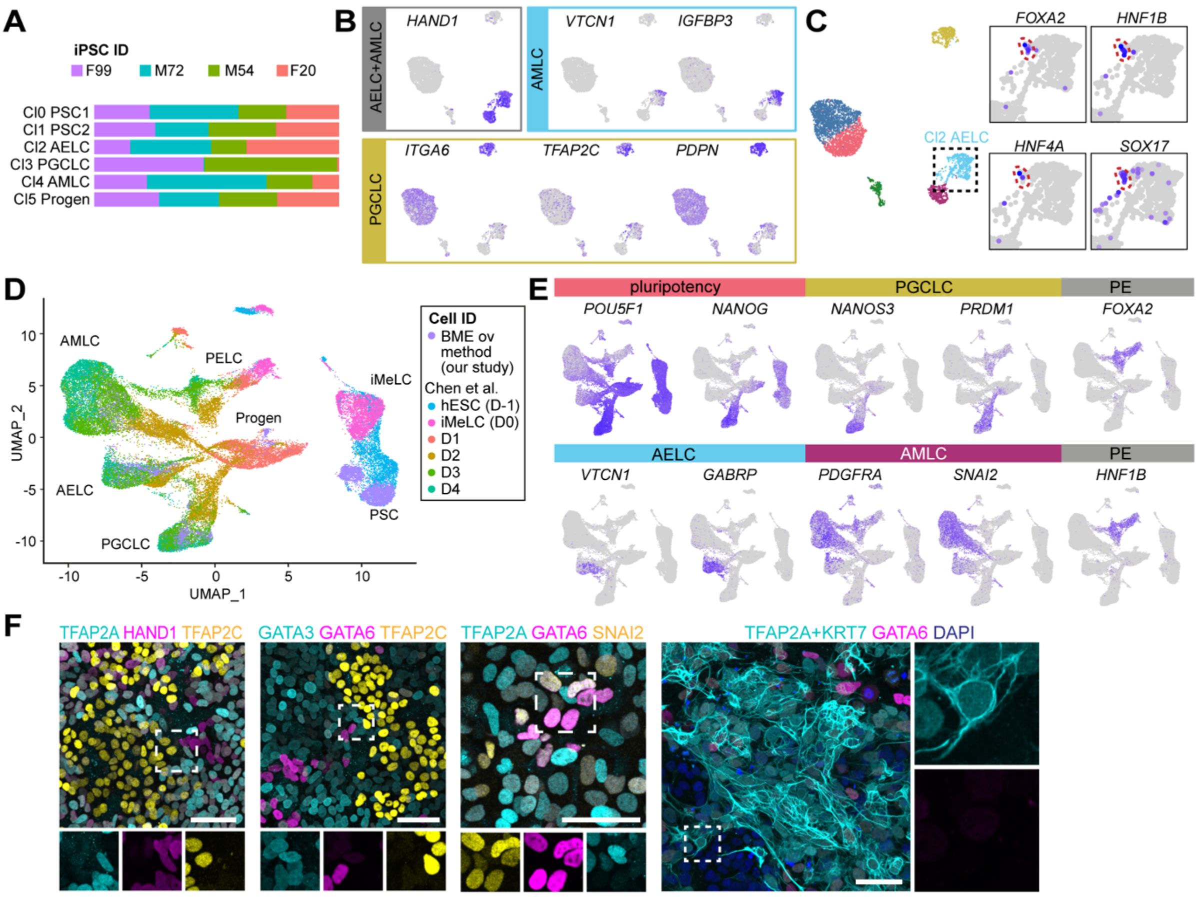


**Figure S2. RNAseq comparing BME overlay method and EB method and verification of markers by IF**

**(A)** Bar graph depicting the contribution per cluster from each cell line.

**(B)** Expression of additional signature genes of cell types of interest on the UMAP plot from Figure 2A.

**(C)** Expression of signature genes of endoderm on the UMAP plot from Figure 2A (close up).

**(D)** UMAP showing integrated the single-cell transcriptomics data from EB-differentiation method (UCLA2 from Chen et al., 2019) and BME overlay method.

**(E)** Expression of additional signature genes of cell types of interest on the UMAP plot from Figure 2D.

**(F)** Immunofluorescence for TFAP2A, HAND1 and TFAP2C; GATA3, GATA6 and TFAP2C; TFAP2A, GATA6 and SNAI2; and TFAP2A+KRT7 and GATA6 at D5 with BME overlay. The dashed box is magnified showing separated channels. Scalebars: 50µm

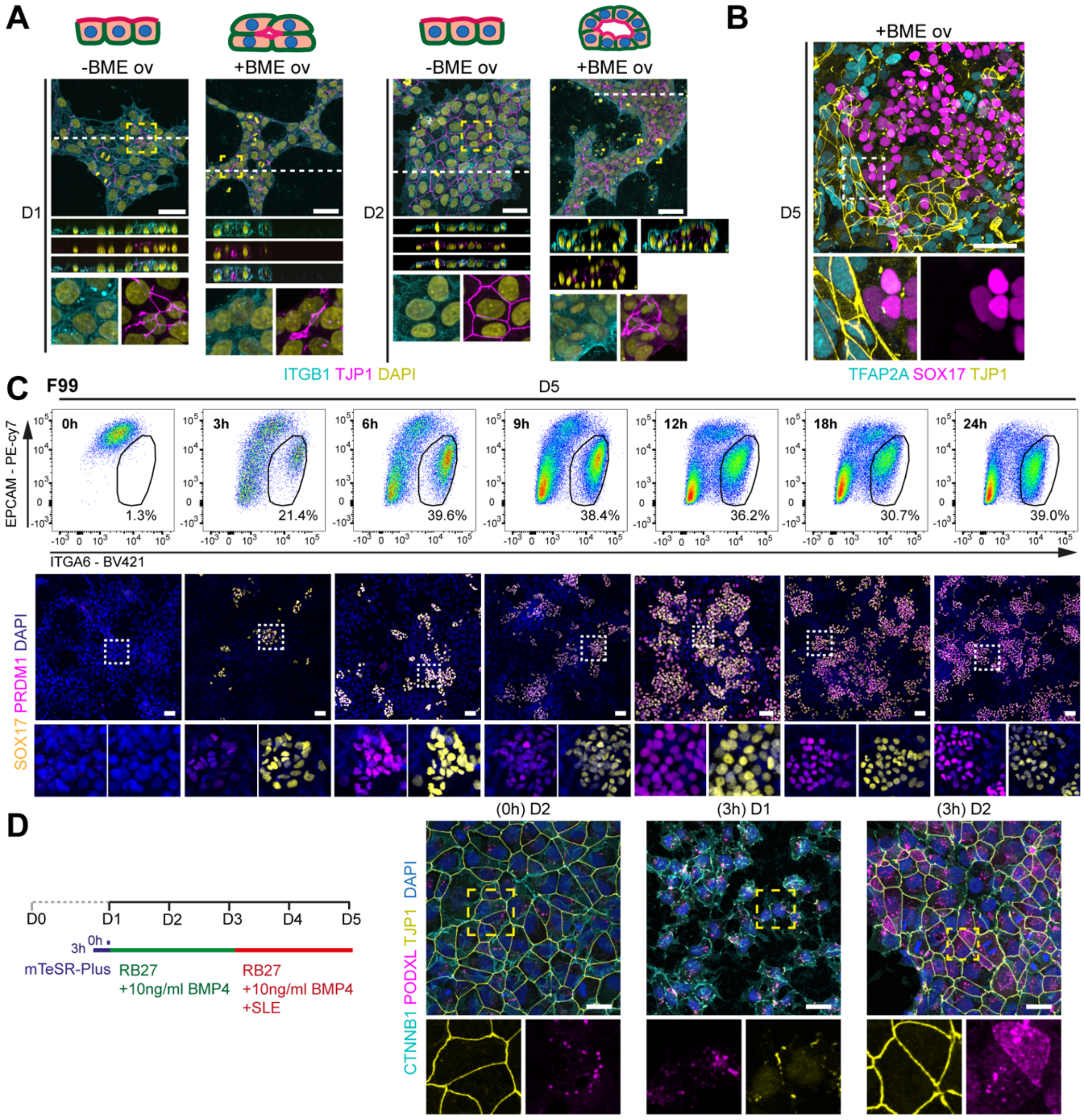


**Figure S3. Lumenogenesis during hPGCLC differentiation with BME overlay**

**(A)** Immunofluorescence for ITGB1 and TJP1 at D1 and D2 with or without BME overlay. White dashed line shows the level of the digital cross section (middle panels) and yellow dashed box is magnified (bottom) showing separated channels. Scale bars: 30µm.

**(B)** Immunofluorescence for TFAP2A, SOX17 and TJP1 at D5 with BME overlay. Dashed box is magnified (bottom) showing separated channels. Scale bars: 50µm.

**(C)** FACS plots depicting the percentage of double EPCAM+ITGA6+ cells at D5 in line F99 to test different priming periods (top) and associated immunofluorescence for SOX17 and PRDM1. Dashed box is magnified (bottom) showing separated channels. Scale bars: 50µm.

**(D)** Experimental scheme depicting the different conditions tested in the absence of BME overlay (left) and immunofluorescence for CTNNB1, PODXL, TJP1 at D1 and D2 to test two different priming periods (right). Dashed box is magnified (bottom) showing separated channels. Scale bars: 50µm.

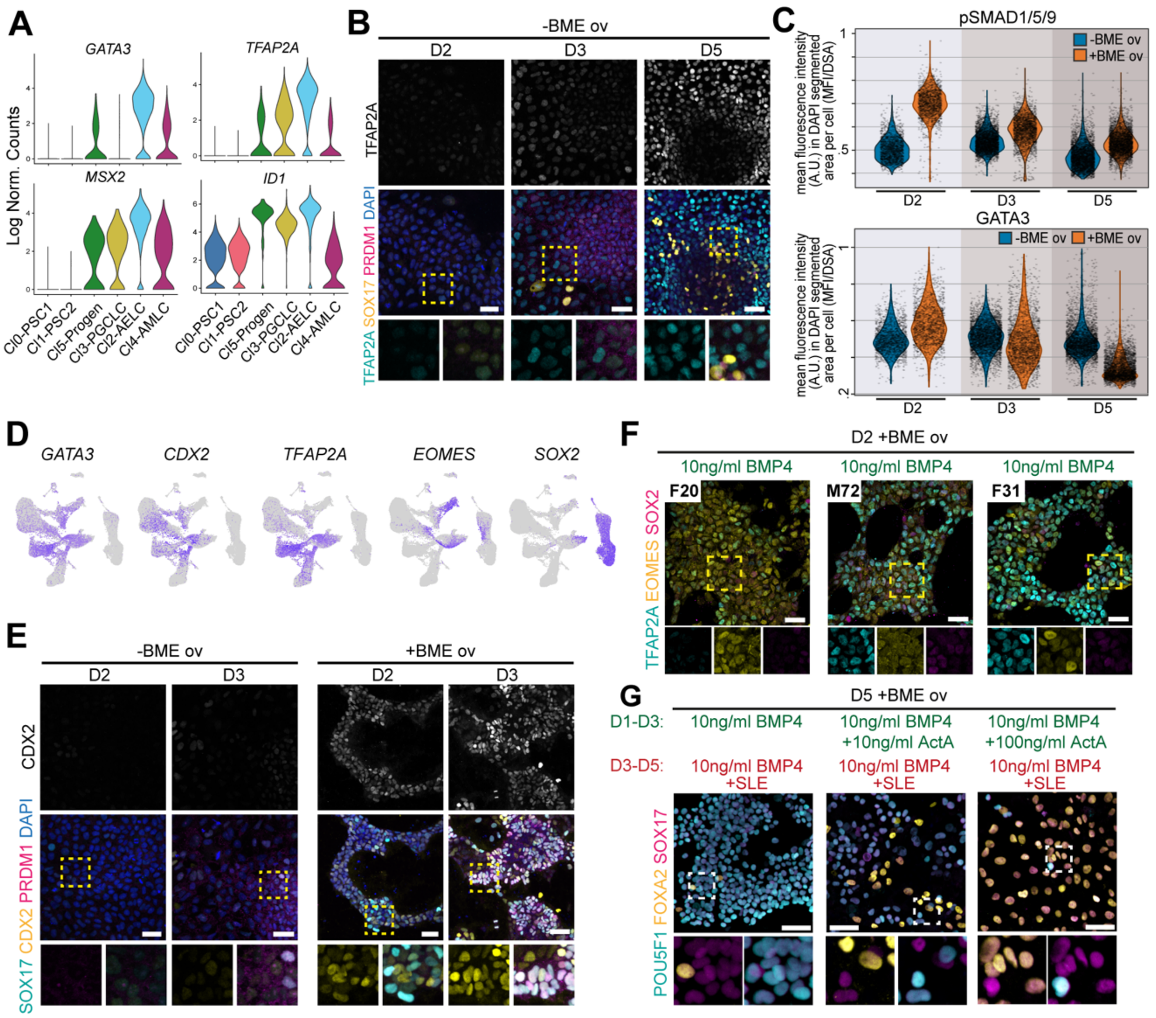


**Figure S4. Characterization of D2-progenitors during hPGCLC differentiation with BME overlay**

**(A)** Violin plots showing expression levels of selected genes of interest per cluster.

**(B)** Immunofluorescence for TFAP2A, SOX17, PRDM1 at D2, D3 and D5 without BME overlay in line M54. TFAP2A is showed on top as single channel. Dashed box is magnified (bellow) showing separated channels. Scale bars: 50μm.

**(C)** Violin plots depict the quantification of the images in Figure 4C as the mean fluorescence intensity in arbitrary units (A.U.) of pSMAD1/5/9 (top) and GATA3 (bottom) in DAPI segmented areas (normalized to 1) per cell.

**(D)** Expression of signature genes of hPGCLC-progenitors on the UMAP plot from Figure 2D.

**(E)** Immunofluorescence for SOX17, CDX2, PRDM1 at D2 and D3 with or without BME overlay. CDX2 is showed on top as single channel. Dashed box is magnified (bellow) showing separated channels. Scale bars: 50μm.

**(F)** Immunofluorescence for TFAP2A, EOMES, SOX2 at D2 with BME overlay in line F20, M72 and F31. Dashed box is magnified (bellow) showing separated channels. Scale bars: 50μm.

**(G)** Immunofluorescence for POU5F1, FOXA2, SOX17 at D5 with BME overlay in the indicated culture conditions. Dashed box is magnified (bellow) showing separated channels. Scale bars: 50μm.
